# A Sister of PIN1 gene in tomato *(Solanum lycopersicum)* defines organ initiation patterns by maintaining epidermal auxin flux

**DOI:** 10.1101/042150

**Authors:** Ciera C. Martinez, Daniel Koenig, Daniel H. Chitwood, Neelima R. Sinha

## Abstract

The spatiotemporal localization of the plant hormone auxin acts as a positional cue during early leaf and flower organogenesis. One of the main contributors to auxin localization is the auxin efflux carrier PIN-FORMED1 (PIN1). Phylogenetic analysis has revealed that PIN1 genes are split into two sister clades; *PIN1* and the relatively uncharacterized *Sister-Of-PIN1 (SoPIN1)*. In this paper we identify *entire-2* as a loss-of-function *SlSoPIN1a* (Solyc10g078370) mutant in *Solanum lycopersicum*. The *entire-2* plants are unable to specify proper leaf initiation leading to a frequent switch from the wild type spiral phyllotactic pattern to distichous and decussate patterns. Leaves in *entire-2* are large and less complex and the leaflets display spatial deformities in lamina expansion, vascular development, and margin specification. During sympodial growth in *entire-2* the specification of organ position and identity is greatly affected resulting in variable branching patterns on the main sympodial and inflorescence axes. To understand how *SlSoPIN1a* functions in establishing proper auxin maxima we used the auxin signaling reporter DR5::Venus to visualize differences in auxin localization between *entire-2* and wild type. DR5::Venus visualization shows a widening of auxin localization which spreads to subepidermal tissue layers during early leaf and flower organogenesis, showing that *SoPIN1* functions to focus auxin signaling to the epidermal layer. The striking spatial deformities observed in *entire-2* help provide a mechanistic framework for explaining the function of the *SoPIN1* clade in angiosperm species.

**Author Summary:** The plant hormone auxin acts as a positional signal in most plant developmental processes. The *PIN-FORMED* family of auxin transporters are the main contributors to auxin localization, especially *PIN-FORMED1*, which has been studied extensively in plant model species *Arabidopsis thaliana*. Members of the PIN-FORMED gene family have been found in all plant species, but there is a scarcity of mutants described outside *Arabidopsis thaliana*. Using *Solanum lycopersicum* (tomato) as a system, this study identifies a loss of function mutant from the *Sister-Of-PIN1* clade in the *SlSoPIN1a* gene. The characterization of this mutant reveals the role of *SlSoPIN1a* in establishing position of organ initiation during shoot and reproductive development, including a role in establishing proper spiral phyllotaxy. We use an auxin visualization technique to conclude *SlSoPIN1a* functions in specifying auxin presence in proper cell layers to establish organ and tissue positioning. This work gives further evolutionary context to how *PIN-FORMED* genes act to establish organogenesis in the plant kingdom.

## Introduction

In plants, cell fate and subsequent tissue formation are mainly determined by positional information rather than cell lineage. The plant hormone auxin acts as a positional cue for proper patterning in many developmental processes, including embryogenesis [1,2], leaf and leaflet initiation [2-6], vascular patterning [5,7-9], root organogenesis [10] and flower initiation [3,11,12]. Auxin presence guides the organization of these processes by inducing changes in transcriptional responses and by affecting cell wall physical properties. The multifaceted role of auxin necessitates a coordinated regulation of auxin influx and efflux carriers which guide auxin transport in a polar fashion, and make up the Polar Auxin Transport (PAT) network.

PAT facilitates auxin action to be precisely coordinated in both a localized and concentration dependent manner. Unlike other known plant hormones, auxin is actively transported in a directional fashion, allowing the creation of spatio-temporally regulated auxin concentrations. The largest contributors of directional transport in the PAT system are the PIN-FORMED (PIN) auxin transporters [13-16]. Most PINs (PIN1, PIN2, PIN3, PIN4 and PIN7) accomplish directional transport by localizing asymmetrically on the plasma membrane of a cell [17], transporting auxin out of the cell in the direction of PIN localization. Auxin, as a weak acid, is freely taken up into a cell, therefore transport of auxin out of the cell by PIN proteins is the determining factor for directional auxin movement [17-19]. The cumulative effect of PIN localization at the tissue level is the establishment of an auxin concentration gradient across a developing organ, and generation of small regions of high auxin concentration called auxin maxima [20-22].

Our understanding *of PIN-FORMED (PIN1)* contribution to plant patterning began with the characterization of the *Arabidopsis thaliana (A. thaliana) pin1 (atpin1)* loss of function mutant [13,26,27]. The defining phenotype of *atpin1* is the presence of radialized “pin-like” structures that are unable to make lateral organs. The *atpin1* phenotype is a consequence of the mutant plants being unable to form auxin maxima required to specify and initiate lateral organs on the flanks of the inflorescence meristem [13,26]. The influence of *AtPIN1* on *A. thaliana* shoot organogenesis varies with developmental age, as the mutant only loses the ability to initiate organs after the floral transition. Prior to reproduction, leaves form on mutant plants, although there are spatial organization problems including aberrant phyllotactic patterning [13,26,27] and leaf and vein developmental abnormalities. These abnormalities, which increase in severity with each developmental stage [27], clearly illustrate that *AtPIN1* contributes to organ establishment during development.

Reiteration of plant modules is a unifying theme during plant development and understanding the formation of these reiterative patterns, especially phyllotaxy, has sparked multidisciplinary interest throughout history. The first molecular marker of leaf organ formation, and thus phyllotactic patterning, is PIN1 localization on the periphery of apical meristems which creates an auxin maximum, marking the site of leaf initiation [3,11,12]. PIN1 predominantly localizes on the L1 (epidermal) layer directing auxin to convergence points, where an auxin maxima is formed and then auxin subsequently becomes directed subepidermally at the site of leaf initiation [4,12]. The transport of auxin through the center of a newly developing leaf continues as the tip of the leaf begins synthesizing auxin, further directing vascular tissue differentiation in its wake [5,7-9]. This pattern of epidermal PIN1 convergence creating subepidermal auxin localization is reiterated throughout development creating many plant reiterative processes including generation of the midvein of leaves [4,12] and leaflets [6], higher-order veins [23], margin development to form leaf serrations [9,24,25] and floral organs specification [3,11,12].

The importance of auxin transport by the PIN transporters in plant development is evidenced by the prevalence of *PIN* genes across the plant kingdom. *PIN* genes have been found in every plant species sampled and in the algal lineage where terrestrial plants emerged, Charophyta [28-30]. In light of their importance in most developmental processes, *PIN* genes have been described as one of the most important gene families guiding plant developmental evolution and plant colonization on land [29,31-35]. Unfortunately there are only a handful of studies that characterize *PIN* gene function outside the model species *A. thaliana*. Recent phylogenetic analysis of *PIN* genes has revealed that most angiosperm species have multiple orthologs of *AtPIN1*, and recent work is in agreement that *A. thaliana* is rare amongst Angiosperm species, in that the Brassicaceae family has likely recently lost a representative in the *“Sister of PIN1 clade” (SoPIN1)* [36-38]. Conservation of PIN-regulated developmental modules is likely species-specific and the extent of divergence in these modules needs to be addressed by analysis of PIN gene function in other species.

*S. lycopersicum* is a model system for studying shoot organogenesis, owing to the large and easily accessible apical meristem and sympodial mode of branching at the floral transition. *S. lycopersicum* has also been used specifically to understand auxin directed developmental mechanisms such as *SlSoPIN1* protein localization in developing organs [6,39-41] and effects of auxin application on organogenesis [3,4,6]. Although *S. lycopersicum* is used extensively as a model system for understanding auxin directed development there is little functional work on *PIN* genes within this species. RNAi knock-down experiments are difficult owing to sequence similarity of the target sequences and have yielded limited insight into *PIN1* function in *S. lycopersicum [42]*. There are 10 *PIN* genes in *S. lycopersicum*, three of which reside in a highly supported phylogenetic clade with *AtPIN1* [36,37,42,43]

To determine *PIN1* function in a broader evolutionary context, we analyzed the function of *SoPIN1* outside the limited context of the Brassicaceae member *A. thaliana*. This study identifies *entire-2*, a previously uncharacterized *SlSoPIN1a* loss of function mutant in *S. lycopersicum*. Phenotypic characterization revealed the role of *SlSoPIN1a* in spatial organization during organogenesis and in leaf, flower, and fruit development. Auxin maxima and auxin-induced gene activity were visualized using an auxin inducible promoter-reporter system, DR5::Venus, and showed that *SlSoPIN1a* loss of function causes an excess of auxin in the apical, inflorescence, and floral meristems and at sites of formation of vasculature causing aberrant developmental responses. We conclude that *SlSoPIN1* regulates auxin patterning by allowing auxin movement in tissue specific cell layers to create a correct spatiotemporal pattern of auxin concentrations needed to guide organ initiation and subsequent morphogenetic processes.

## Results

### There was a *SoPIN1* gene duplication event prior to the diversification of the Solanaceae

In *S. lycopersicum*, there is one true *AtPIN1* ortholog, *SlPIN1* (Solyc03g118740), and two *SoPIN1* genes, *SlSoPIN1a* (Solyc10g078370) and *SlSoPIN1b* (Solyc10g080880) (Figure 1) [42,43]. Previous phylogenetic analysis places both *SlSoPIN1a* and *SlSoPIN1b* genes together on a single branch tip, suggesting a recent *SoPIN1* gene duplication event in the branch leading to *S. lycopersicum [36,37]*. These reports suggested that a duplication event in the *SoPIN1* clade in *S. lycopersicum* occurred roughly sometime after the divergence between *S. lycopersicum* and *Mimulas guttatus [36,37]*. To gain a more precise understanding of the history of the *SoPIN1* clade, we performed phylogenetic analysis on *PIN1* and *SoPIN1* genes sampling Solanaceae more extensively by including *Capsicum annuum, Nicotiana benthamiana, Solanum habrochaites, Solanum lycopersicum, Solanum pennellii, Solanum pimpinellifolium*, and *Solanum tuberosum*. In addition we included seven other representative Eudicot species *(Arabidopsis thaliana, Citrus clementina, Capsella rubella, Cucumis sativus, Medicago truncatula*, and *Mimulus guttatus)* (Figure 1A), and used *A. thaliana PIN3* as an outgroup. We performed phylogenetic analysis using the RAxML maximum likelihood method [44] with orthologous cDNA sequences of *SoPIN1* and *SlPIN1* genes. Bootstrap analysis was performed on 1,000 runs to obtain statistical confidence.

**Figure 1.**
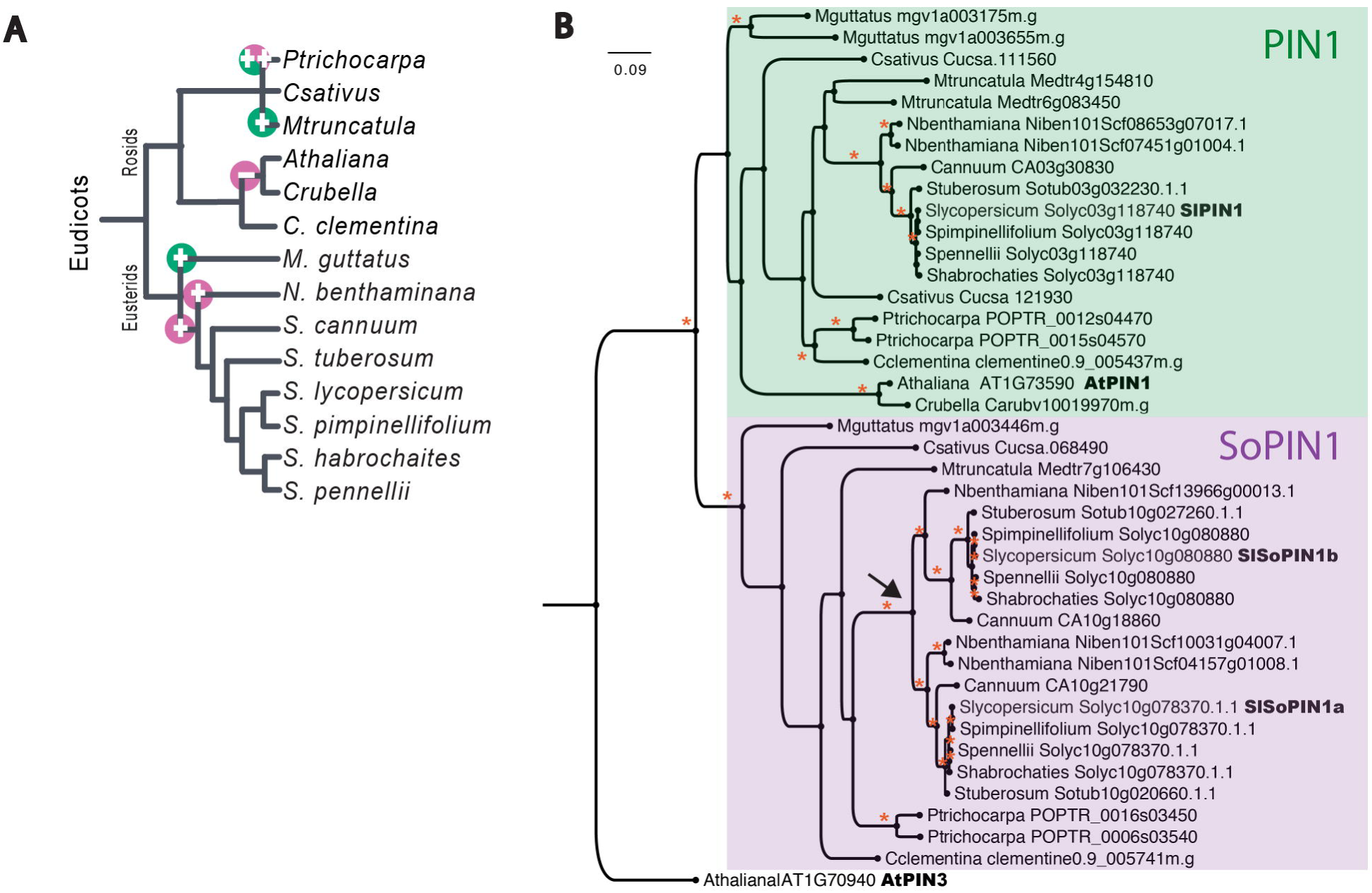
Phylogenetic analysis of dicot *SoPIN1* and *PIN1* genes. (A) Phylogenetic tree showing relatedness of species sampled in this analysis. Major gene duplication (+) and gene loss events (-) in *PIN1* (green) and *SoPIN1* (pink) sequence evolution inferred from phylogenetic tree shown in (B). (B) Maximum likelihood phylogenetic tree of *PIN1* and *SoPIN1* gene divergence. Black arrow shown in (B) infer *SoPIN1* gene duplication event in Solanaceae *SoPIN1* genes. Species names were abbreviated as follows *Arabidopsis thaliana (Athaliana), Capsicum annuum (Cannuum), Citrus Clementina (Cclementina), Capsella rubella (Crubella), Cucumis sativus (Csativus), Medicago truncatula (Mtruncatula), Mimulus guttatus (Mguttatus), Nicotiana benthamiana (Nbenthamiana), Solanum habrochaites (Shabrochaites), Solanum lycopersicum (Slycopersicum), Solanum pennellii (Spennellii), Solanum pimpinellifolium (Spimpinellifolium)*, and *Solanum tuberosum (Stuberosum). \** All nodes have at least 95% bootstrap support. Scale represents 0.09 substitutions per site.

*PIN1* genes fall into two highly supported sister clades, as shown in previous work [36,37], one group which we will continue to refer to as PIN1, where *AtPIN1* (AT1G73590) resides, and a second clade, *Sister-of-PIN1 (SoPIN1)* (Figure 1B). All species sampled have at least one gene represented in each of the *PIN1* and *SoPIN1* clades, with the exception of *A. thaliana* and the closely related species *C. rubella* (Figure 1A and B), demonstrating a likely loss of the *SoPIN1* clade in Brassicaceae, as recently reported [36,37]. Within the *SoPIN1* clade, Solanaceae *SoPIN1* genes are split into two clear groups - each with at least one member from each Solanaceae species sampled (Figure 1B). All other Eudicot species have genes that fall outside these two Solanaceae specific groups, suggesting the *SoPIN1* duplication event occurred just prior to Solanaceae speciation. The function of these duplicated *SoPIN1* genes has never been explored explicitly.

### *e-2* has a stop codon in *SlSoPIN1a*

To compare the function of *SoPIN1* and *PIN1* genes in Solanaceae to the function of *PIN1* in *A. thaliana*, we screened the Tomato Genetics Research Center (TGRC) mutant database for candidate *pin1* and *sopin1* mutants in *S. lycopersicum*. Guided by the phenotypic characterization of *A. thaliana* and *M. truncatula pin1* mutants, we searched for monogenic mutant lines located in the approximate genomic regions of the *S. lycopersicum SlPIN1, SlSoPIN1* and *SlPIN1* genes that also displayed leaf phenotypes. In close proximity to *SlPIN1* on Chromosome 3, there are two mutant lines which display aberrant leaf phenotypes, *divaricata (div)* and *solanifolia (sf)*. Both of the *SoPIN1* genes, *SlSoPIN1a* and *SlSoPIN1b* reside in close proximity to each other on Chromosome 10, along with three leaf phenotype mutants: *oivacea (oli), entire-2 (e-2)*, and *restricta (res)*. We grew all five mutant lines and scored them for spatial arrangement phenotypes in shoot and leaf morphology similar to phenotypes of *A. thaliana* and *M. truncatula sopin1 (slm)* mutants [13,27,45]. Of the mutant lines, *sf, e-2*, and *div* were characterized as having leaf phenotypes (S1 Figure), but only *e-2* and *sf* possessed abnormalities including aberrant vasculature, fused cotyledons, and laminar tissue similar to what was measured in the *M. truncatula sopin1* mutant and *atpin1* mutant lines [13,27,45]. Next we compared the nucleotide sequences of all three *PIN1 genes* in *sf, e-2*, and *div* mutant lines and found that all sequences were identical to that in their wild type background with the exception of *e-2*. The *SlSoPIN1* sequence in *e-2* plants harbors a single nucleotide change from C to T at position 490 (C-T^490^) causing a premature stop codon in the translated amino acid sequence (Figure 2A), suggesting *e-2* as a candidate *sopin1* loss of function mutant. Organ patterning varied considerably between *e-2* individuals, and the most consistent phenotype was a deviation from the wild type spiral phyllotactic pattern. This aspect of the phenotype was subsequently used for further co-segregation and complementation analyses.

**Figure 2.**
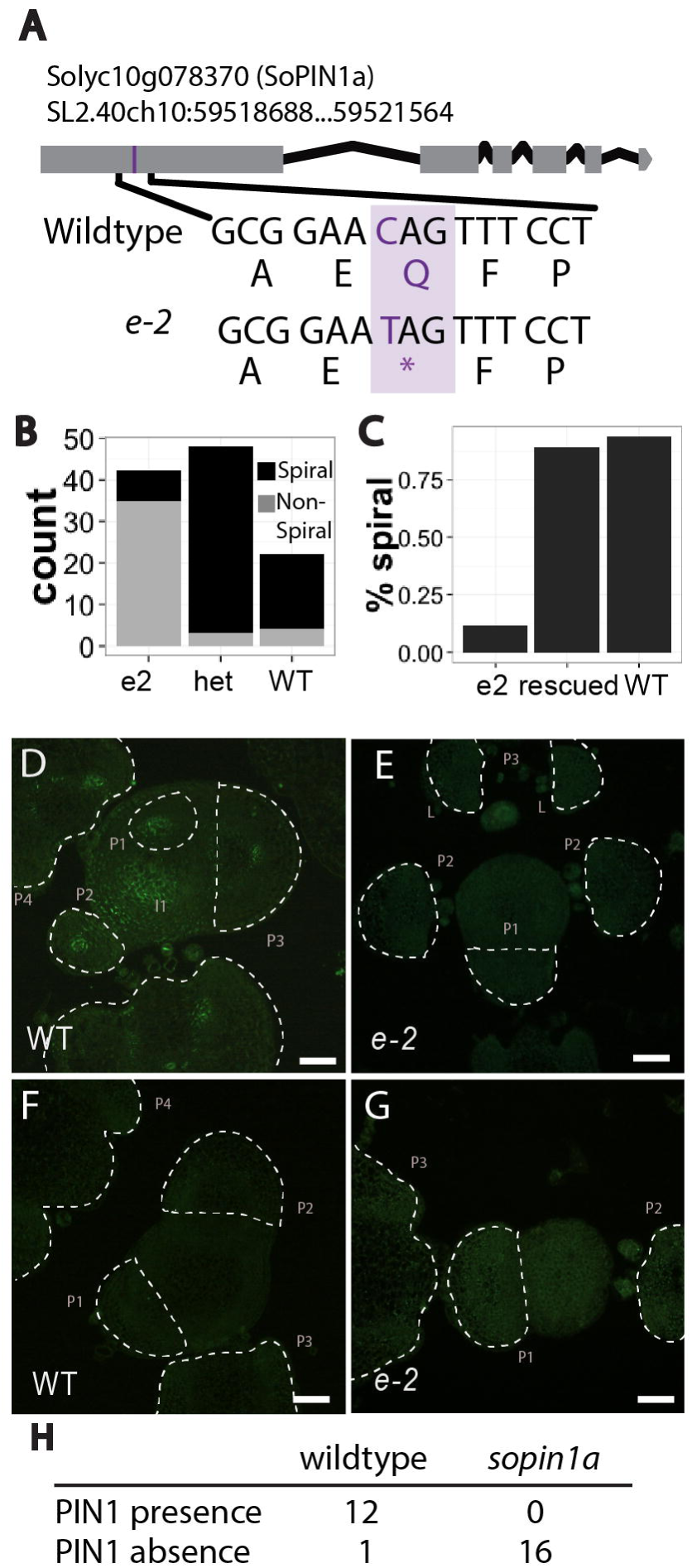
The *e-2* phenotype is caused by lack of *SlSoPIN1a* function. (A) Map illustrating location of nucleotide change (C-T^490^; purple) in the *e-2 SlSoPIN1a* gene which results in a premature stop codon (*) in the translated amino acid sequence. (B) Results of a co-segregation experiment showing the deviation from wild type spiral phyllotaxy segregates with the homozygous C-T^490^ polymorphism in the progeny genotyped. (C) Bar plot illustrating that spiral phyllotaxy segregates with homozygous C-T^490^ individuals rescued with a functional *PIN1* gene (AtpPIN1::PIN1::GFP). Rescued individuals are those homozygous for C-T^490^ and genotyped as having AtpPIN1::PIN1::GFP presence. (D)-(G) Immunolocalization experiments (E) and (F) showing SlSoPIN1a antibody signal (green) is only found in wild type (E) apices and SlSoPIN1a protein was never observed in *enitre-2* (E) apices. (G) and (G) no signal was found in control using only secondary antibody without primary SlSoPIN1a anti-body. (I) Table summarizing immunolocalization results. (D)-(G) Scale bars = .1 mm.

### C-T^490^ co-segregates with the *e-2* phenotype

We tested if C-T^490^ was responsible for the *e-2* phenotype by asking if C-T^490^ co-segregates with the phyllotaxy phenotype. We crossed *e-2* with the wild type background, self-pollinated the heterozygous progeny, and scored for presence and absence of the spiral phyllotactic pattern in the F2 population (n = 112). Plants were phenotyped 30 days after germination, when Leaf 1 to ~Leaf 7 are clearly visible. Of the individuals homozygous for C-T490 86.4% scored as having an absence of a spiral phyllotaxy. In the parental *e-2* homozygous population 88.2% of the individuals show aberrant phyllotaxy (Figure 2B). In the segregating F2 population 81.8% individuals homozygous for the wild type allele possessed spiral phyllotaxy, while 93.7% of the heterozygotes exhibited this trait. Assuming plants heterozygous and homozygous for the wild type allele display a spiral phyllotactic pattern 100% of the time, the segregation of the phyllotaxy phenotype did not differ from expected based on genotype (chi-squared test, p-value = 1.00). These results indicate that the C-T^490^ SNP in *e-2* is a likely candidate for the deviation in spiral phyllotaxy seen in *e-2* and further supports our hypothesis that *e-2* is a *slsopin1a* mutant.

### *e-2* phenotype can be complemented with a functional *PIN1* gene

To further establish that the *e-2* phenotype is caused by a lack of *SlSoPIN1a* function, we asked if a functional PIN1 protein could rescue the phyllotaxy phenotype in *e-2*. Since SoPIN1 is absent in *A. thaliana*, we tested if a functional copy of *AtPIN1* is able to rescue SoPIN1 protein function, as seen previously with a similar construct that rescued the *mtsopin1* mutant phenotype [12,45]. We evaluated the presence or absence of spiral phyllotaxy in a population (n = 117) homozygous for C-T^490^ and segregating for pPIN1::PIN1::GFP, a construct containing a functional *PIN1* gene and promoter from *A. thaliana* [11,39]. All individuals of the population were genotyped as homozygous for C-T^490^ and 72.22% of the population genotyped positive for the pPIN1::PIN1::GFP construct. If pPIN1::PIN1::GFP is capable of rescue, we would expect approximately 72.22% of the population to have a spiral phyllotactic pattern. We found 77.16% of the population displayed the expected rescued spiral phyllotactic phenotype, which is not significantly different from expected (chi-squared test, p-value = 0.3149). Most individuals genotyped for pPIN1::PIN1::GFP (88%) displayed the wild type spiral phenotype (Figure 2C). Further, leaf divergence angle in complemented individuals showed a similar angle of divergence to that found in WT (S2 Figure E). Thus a functional *PIN1* is capable of rescuing the aberrant phyllotaxy found in *e-2* individuals homozygous for C-T^490^.

The phenotypic characterization of *e-2* revealed the role of *SlSoPIN1a* in spatial organization during leaf and inflorescence development. We further asked if there is a difference in auxin signaling and localization in *e-2* that could explain these phenotypes. To verify the role of *PIN1* in the developmental processes characterized above we used the *A. thaliana* pPIN1::PIN1::GFP reporter line, which showed SlSoPIN1 protein localization in tomato highly similar to that seen in *A. thaliana* [39]. Since pPIN1::PIN1::GFP complements the *e-2* phenotype, as expected, we observed no difference in pPIN1::PIN1::GFP expression in complemented *e-2* lines (S3 Figure A-H).

### C-T^490^ causes loss of *SoSlPIN1* function in *e-2*

We next asked how C-T^490^ affects gene function at the transcriptional and translational level. Using primers positioned either before or after the C-T^490^ in *slsopin1* individuals, we found that the transcript was present and levels were not significantly different between the *e-2* and wild type backgrounds (S2 Figure C), therefore C-T^490^ does not appear to affect transcription levels in *e-2*. Differences at the translational level were assessed by the presence or absence of the SlSoPIN1a protein using an antibody raised against the *SlSoPIN1a* sequence [39]. In wild type, we observed SlSoPIN1a protein localization in incipient primordia and provascular tissue (Figure 2D; S3 Figure I-P) as observed previously [39]. In contrast, SlSoPIN1a protein was always absent in *e-2* apices (Figure 2E and G). In all, we performed immunolocalization on twelve apices of both *e-2* and wild type. SlSoPIN1a protein localization was never observed in *e-2* individuals, while SlSoPIN1a antibody presence was found in all but one WT apex sampled (Figure 2H). These results indicate that C-T^490^ does not affect *SlSoPIN1a* transcription, but prevents accumulation of functional *SlSoPIN1a* by preventing translation in *e-2* apices, further confirming that *e-2* is a loss of function *slsopin1a* mutant.

### *SoSlPIN1a* regulates phyllotaxy

We quantified how *e-2* plants deviate from spiral phyllotactic patterning by measuring the leaf divergence angle in both wild type (n= 114) and *e-2* (n = 124). *S. lycopersicum* spiral phyllotactic patterning generally follows a regular repeating angle of divergence, with leaves emerging around 137.5° [20] (Figure 3A-D). In wild type, we measured divergence angles clustered around 137.5° as expected, while in *e-2*, the divergence angles varied extensively, with measurements clustering in the upper limits of divergence ~ 180° (Figure 3C). Around half of *e-2* individuals (44.31%) display a distichous or decussate phyllotaxy, as leaves alternatively emergence at around 180° and 90° (S2 Figure E). There are few known plant mutants, which switch phyllotactic pattern, but in three characterized phyllotaxy mutants meristem size has been attributed as a possible determining factor [46-48]. The spatial constraints of meristem size influence where new auxin maxima can form to initiate new leaves. In order to assess if meristem size in *e-2* is a contributing factor to the *e-2* phyllotaxy phenotype, we measured the width and height of wild type (n = 34) and *e-2* (n = 46) meristems (Figure 3G and H). Overall *e-2* meristems are significantly larger in height (Welch Two Sample t-test, p-value = 8.729e-06) but have no significant difference in meristem width (Welch Two Sample t-test, p-value = 0.1767) (Figure 3I), suggesting meristem size is one contributing factor to the aberrant phyllotaxy observed in *e-2*.

**Figure 3.**
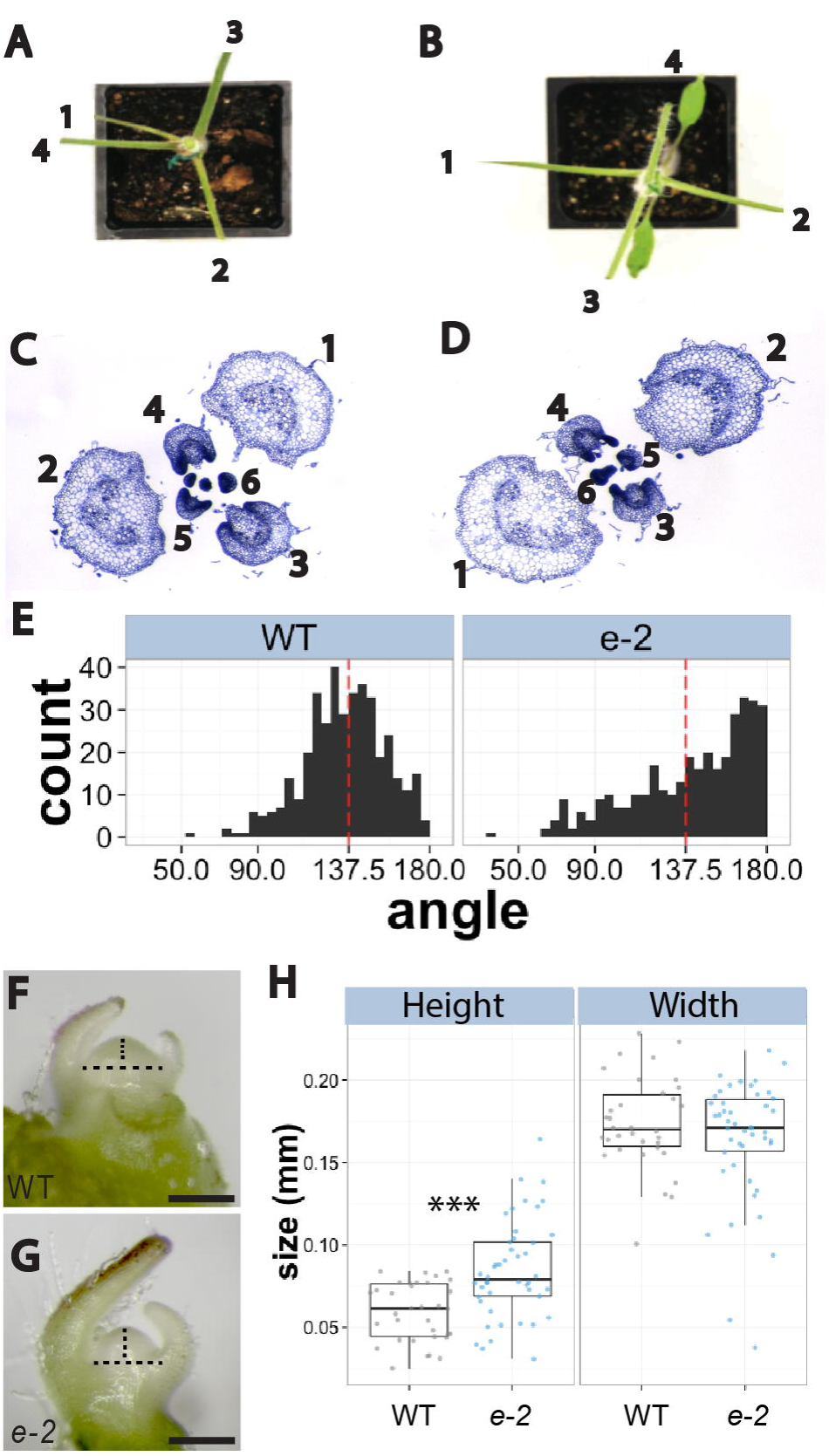
*SoPIN1a* regulates phyllotactic patterning in *S. lycopersicum*. (A) and (C) wild type plants displaying spiral phyllotactic patterning. (B) and (D) *e-2* plants displaying a distichous phyllotaxy. (E) Divergence angles across leaves 1-8. (F) to (H) Photographs of apical meristems from (F) wild type and (G) *e-2* Dotted lines represents how measurements were obtained for length and width in (F) and (G). (H) Differences in meristem size between wild type (grey) and *e-2* (blue). *** indicates p-value < .0005. Scale bars = 0.2 mm in (F) and (G).

### *SoSlPIN1a* regulates spatial identity during leaf development

One of the most striking phenotypes of *e-2* is the improper specification of leaf tissues. During leaf initiation, midvein formation begins as auxin is transported subepidermally through the center of the newly established leaf [4,12]. The midveins of *e-2* leaves are thickened and often indistinguishable from secondary vasculature, giving the appearance of multiple mid veins (S4 Figure A). This trend of thickened vasculature is also displayed in higher order veins, attributed to irregular spacing, resulting in fused veins and increased tracheary elements (Figure 4E-H). This phenotype has also been seen in *atpin1* mutant [27] and can be replicated by application of NPA, an auxin transport inhibitor, resulting in leaves with similar phenotype as seen in *e-2* (S2 Figure F-I) [7,27,49]. Vasculature in the petiole is also affected in *e-2*, as cross sections through the petiole reveal a lack of separation of vascular bundles seen in wild type (S4 Figure F-K). In wild type lamina, blade tissue is smooth and lies flat, while *e-2* leaf tissue shows bulging of intervein tissue that worsens as development proceeds (Figure 4A and B; S2 Figure B-E).

**Figure 4.**
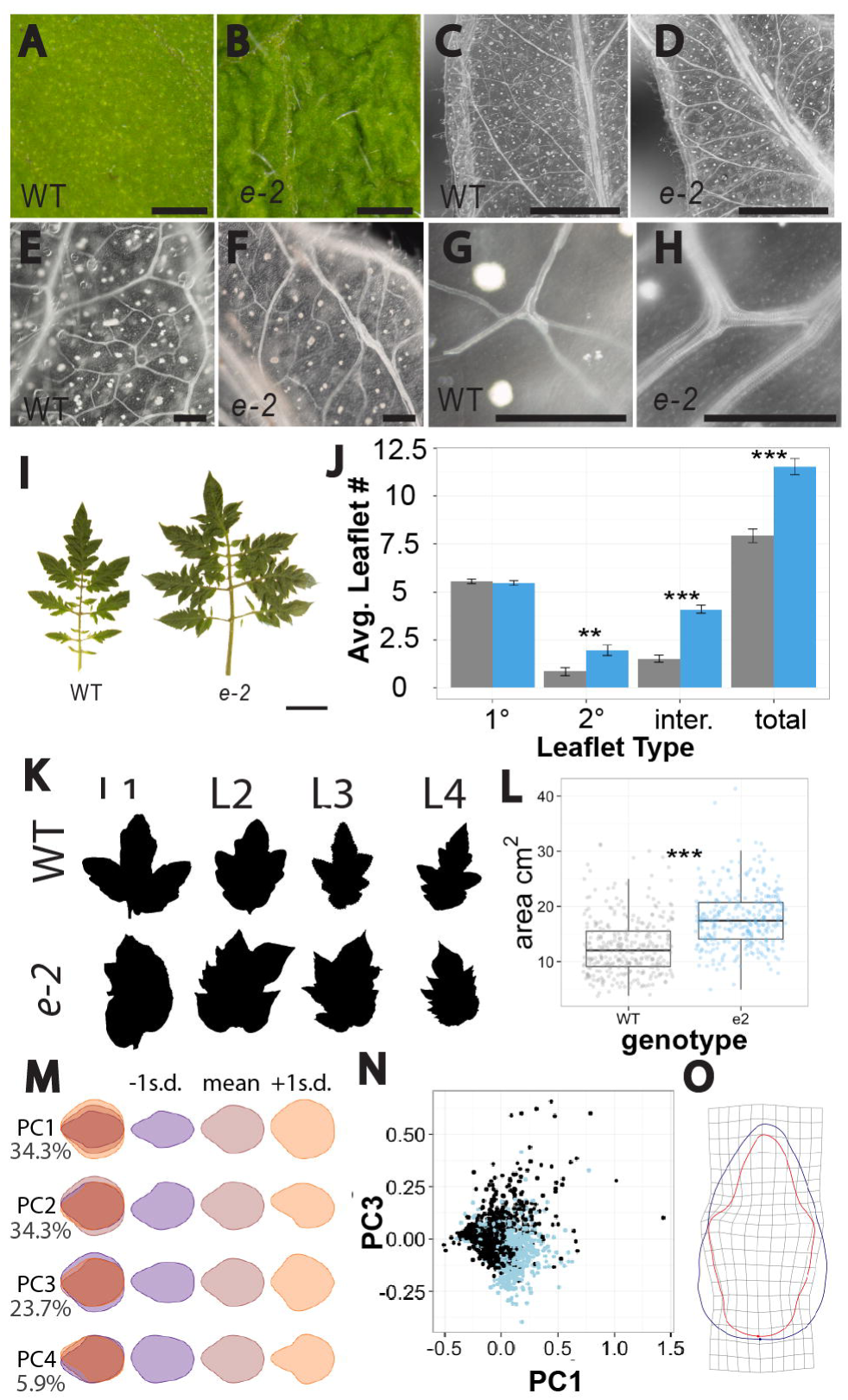
Leaf development in *e-2*. (A) and (B) lamina tissue from Leaf 4 (A) wild type and (B) *e-2* three week old plants. (C) to (H) Leaf 2 from two week old plants cleared with hydrochlorate showing midvein development (C) and (D), vascular tissue near leaf margin (E) and (F), and treachery branching (G) and (H) in wild type (C), (E), and (G) and *e-2* (D),(F),(H). (I) and (J) Leaf 4 of five week old plants showing *e-2* leaves are larger and less complex than wild type. (J) Bar graph illustrating average leaflet number of Leaf 4 leaves from six week old plants, showing *e-2* (blue; n = 70) leaves are significantly less complex compared to wild type (grey; n= 49). Error bars represent Mean ± Standard Error (SE). (K) Representative binary terminal leaflet outlines from four week old plants. (L) Boxplot showing leaflet area from fully expanded leaves 1 and 2 of wild type (grey; n = 316) and *e-2* (blue; n = 315). (M) Principal Components (PC) 1-4 illustrated as leaf outlines which include −1 standard deviations (purple) and +1 standard deviations (orange) along each axis and mean outline (gray). (N) Scatter plot of PC1 and PC3 obtained from elliptical fourier analysis performed on leaf outlines from wild type (black) and *e-2* (blue). (O) Average terminal leaflets outline from Leaves 1-4, derived from elliptical fourier analysis. Welch’s t-test P-value < 0.01 = **, P-value < .0001 = ***. Scale bars = 1 mm in (A) to (D), 0.1 mm in (E) to (H), and 2 cm in (I).

The leaves of *e-2* are significantly less complex due to a reduction of secondary and intercalary leaflets (Welch Two Sample t-test, p = 0.00198 and 4.033e-15, respectively) (Figure 4J). Leaf margins in *e-2* have irregular lobing and sharp serrations compared to wild type (Figure 4K). In order to quantify leaf shape explicitly, shape differences were derived from characterization of the terminal leaflet of mutant and wild type. Overall, *e-2* leaflets have a significantly larger area (Welch Two Sample t-test, p = < 2.20 e-16) (Figure 4L) and have higher circularity, a measure of leaflet serration and lobing (Welch Two Sample t-test, p=2.391e-13), than wild type individuals (S1 Table). We used Elliptical Fourier analysis [50,51] to quantify differences in shape through measurement of leaflet outlines. Using Principal Component Analysis (PCA), we visualized patterns of variance that exist between the *e-2* and wild type lines. There is a clear separation between WT and *e-2* along PC1 explaining leaf width and lobing (Figure 4M and N). PC2 explains asymmetry common to both genotypes (S4 Q and R). The outline shape varied extensively between *e-2* leaflets, as lobing and serrations were placed seemingly randomly (Figure 4K) resulting in the average *e-2* outline as a near circular shape compared to wild type, which usually consists of three prominent lobes (Figure 4O). As seen in other loss of function *sopin1* and *pin1* mutants [1,11,45], lateral organ morphogenesis is often perturbed during cotyledon development in *e-2* (S4 Figure L-O). Thus, *e-2* individuals are capable of initiating leaves with all the typical leaflets and tissues, although placement of these features is highly irregular. Taken together these results suggest *SoSlPIN1a* functions in proper spatial organization during leaf initiation and morphogenetic processes.

### *SoSlPIN1a* also regulates spatial patterning and specification of organ identity during sympodial growth

Studies have shown that *PIN 1*-directed auxin transport is necessary for floral initiation and development [3,11,12]. The role of *PIN1* in floral organ and inflorescence development is particularly difficult to characterize in *A. thaliana*, owing to the lack of inflorescence initiation in *atpin1* mutants [26], therefore we were particularly interested in characterizing *SoPIN1* function in flower development using *e-2*. In *S. lycopersicum*, monopodial growth begins as leaves initiate from a single monopodial meristem (MM). After the initiation of around 7 - 12 leaves at the MM, plants begin sympodial growth [52]. At the start of sympodial growth the MM produces 1. an inflorescence meristem (IM) and 2. a sympodial meristem (SYM) (Figure 5B). Sympodial growth is delayed in *e-2* likely from a slower rate of initiating leaves (Welch Two Sample t-test, p-value = 0.019) (S4 Figure P). In wild type plants, sympodial growth continues on the main axis of the plant, reiterating through distinct sets of sympodial modules. Each sympodial module consists of a single leaf and an inflorescence branch (Figure 5A). The SYM continues growth on the main axis of the plant while the sympodial inflorescence meristem creates an inflorescence branching structure by further sympodial branching to produce determinate floral organs.

**Figure 5.**
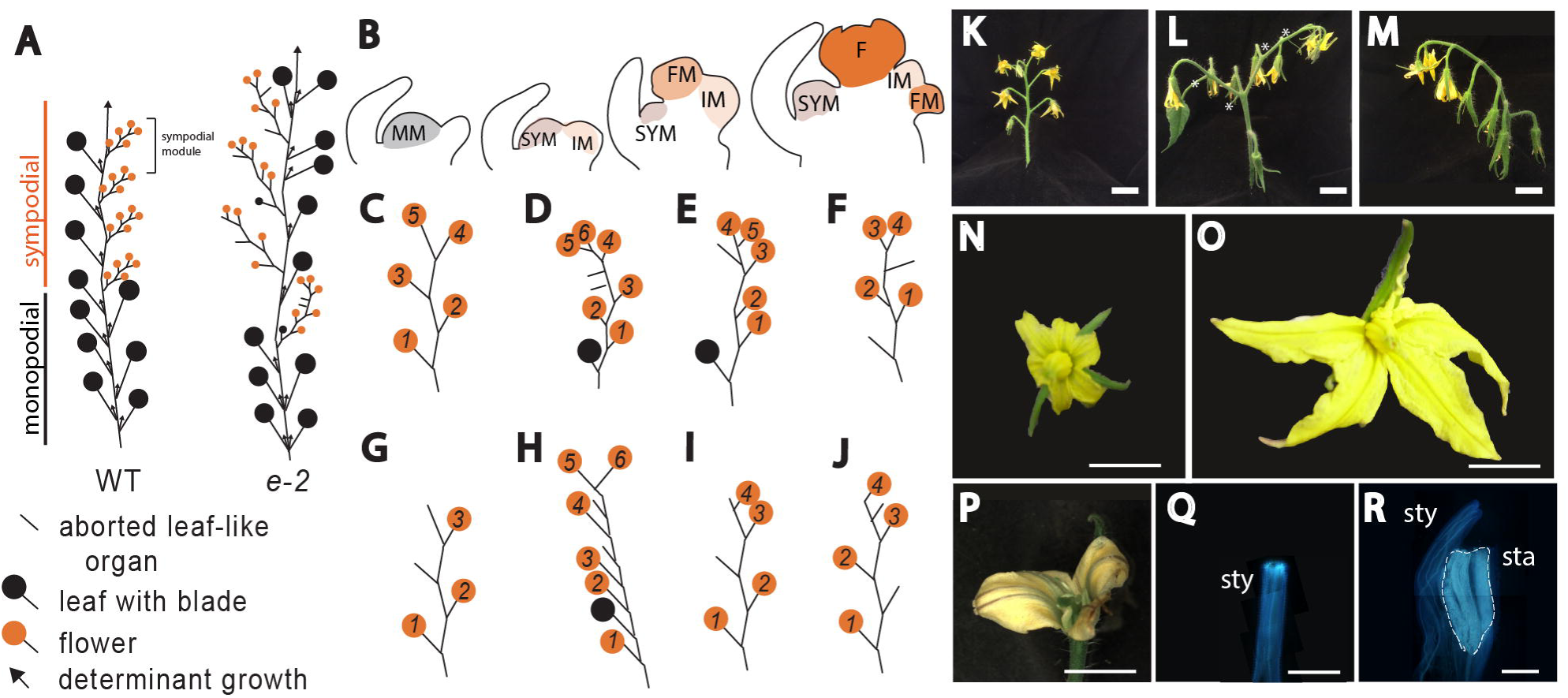
Loss of *SlSoPIN1a* function results in altered organ initiation and morphogenesis during sympodial and flower growth. (A) Schematic example of branching patterns showing inconsistent branching in wild type compared to *e-2*. (B) Schematic illustrating meristem maturation in wild type *S. lycopersicum*. During monopodial growth leaves initiate on the monopodial meristem (MM). Sympodial growth begins as the MM transitions to a sympodial meristem (SYM) and a inflorescence meristem (IM). The IM produces a determinate floral meristem (FM) which terminates as a single flower (F). Sympodial branching continues on the inflorescence as a new IM is initiated. (C) to (J) Schematic examples of inflorescence branching patterns. (J) wild type inflorescence (L) to (M) *e-2* inflorescence. Flower size in wild type (N) is smaller and flowers consistently show five petals, while *e-2* flowers (O) and (P) are often larger having inconsistent petal number due to within whorl tissue fusion events. (P) and (R) Organ fusion events are also observed between wholes, as seen (R) *e-2* as stamens (sta) are fused to style (sty). Wild type style (Q). Scale bars = (K),(L),(M) is 2 cm, (N), (O), (P) is 10mm, and (Q) and (R) is 1mm.

The specification of flower or leaf can occur during sympodial growth on both the main axis and in the inflorescence in some plants. Within Solanaceae, it is not uncommon to see bracts subtending flowers, as seen in *S. pennellii, S. habrochaites*, and *N. benthamiana* (personal observation). However, in *S. lycopersicum* th*e* IM always produces only floral units *[53]*. Surprisingly, in *e-2*, the inflorescence branching pattern is variable creating many variations of the sympodial branching module, including specification of complex leaves, simple leaves, and aborted leaf-like structures in between flower units of the *e-2* inflorescence (Figure 5C-J and K-M; S5 D and I Figure). In *e-2*, both the number of floral units and flower branching structures deviate from the wild type pattern of around five floral units, which normally creates a zig zag branching pattern (Figure 5C and K). We utilized tissue specific gene expression data from early stages of inflorescence [54] to analyze gene expression differences in *SlPIN1, SlSoPIN1a* and *SlSoPIN1b* (S7 Figure). Gene expression of *SlPIN1* remains constant across sympodial inflorescence establishment, the *SoPIN1* genes *(SlSoPIN1a* and *SlSoPIN1b)* have greater variability throughout the different stages and even vary compared to each other, suggesting a possible subfunctionalization during inflorescence establishment. These results, along with *e-2* phenotypic characterization during sympodial growth, suggest *SlSoPIN1a* influences both the architecture of the inflorescence and also the specification of inflorescence organs, possibly by changing gene expression levels during sympodial growth.

Flower differentiation proceeds by the initiation of concentric whorls of lateral organs - the sepals, petals, stamens, and carpels [55]. Plants that are incapable of transporting auxin, either through genetic or chemical disruption, are either unable to form flowers [26] or display aberrant floral organ specification and positioning [26]. In both heterozygous and to a greater extent, homozygous *e-2* individuals, flowers range in deformities from appearing unaffected to extremely disordered, including fusion of flower organs both within whorls (Figure 5M) and between whorls (Figure 5O), loss and gain of organs, and a general increase in organ size (Figure 5L; S5 Figure A-C). Differences in size and shape are continued into fruit development, as *e-2* fruit are often elongated with problems in seed set and placental development (S6 Figure). The fruit and flower phenotypes in *e-2* shows that *SlSoPIN1a* plays a role in reproductive development by regulating organ initiation and size.

### *e-2* shows more diffuse and larger auxin foci during leaf initiation and early leaf development

To visualize potential disruptions in auxin localization in *e-2*, we used an auxin-inducible promoter-reporter system, DR5::Venus [40], often used as an indirect auxin reporter [9,12,37,56-58]. In wild type, during leaf initiation on the MM, DR5::Venus is only found at sites of leaf initiation (P0), with the fluorescence pattern appearing as a wedge shape pointing inward toward the center of the meristem (Figure 6A, C, E, and G). In *e-2*, DR5::Venus signal is seen throughout the MM (Figure 6B and F), suggesting an impairment in auxin transport in the MM. In addition, DR5::Venus fluorescence in *e-2* is found throughout the epidermal layer and persists into the subepidermal layers (Figure 6H). There is no clear demarcation of DR5::Venus signal between P0 and meristem in *e-2*, consistent with a diffuse auxin maximum (Figure 6F and H) as opposed to the wedge shaped one seen in wild type (Figure 6E and G). In addition the DR5::Venus signal at P0 in *e-2* does not penetrate as deep into subdermal layers (Figure 6F and H), again suggesting a defect in auxin transport in the mutant.

**Figure 6.**
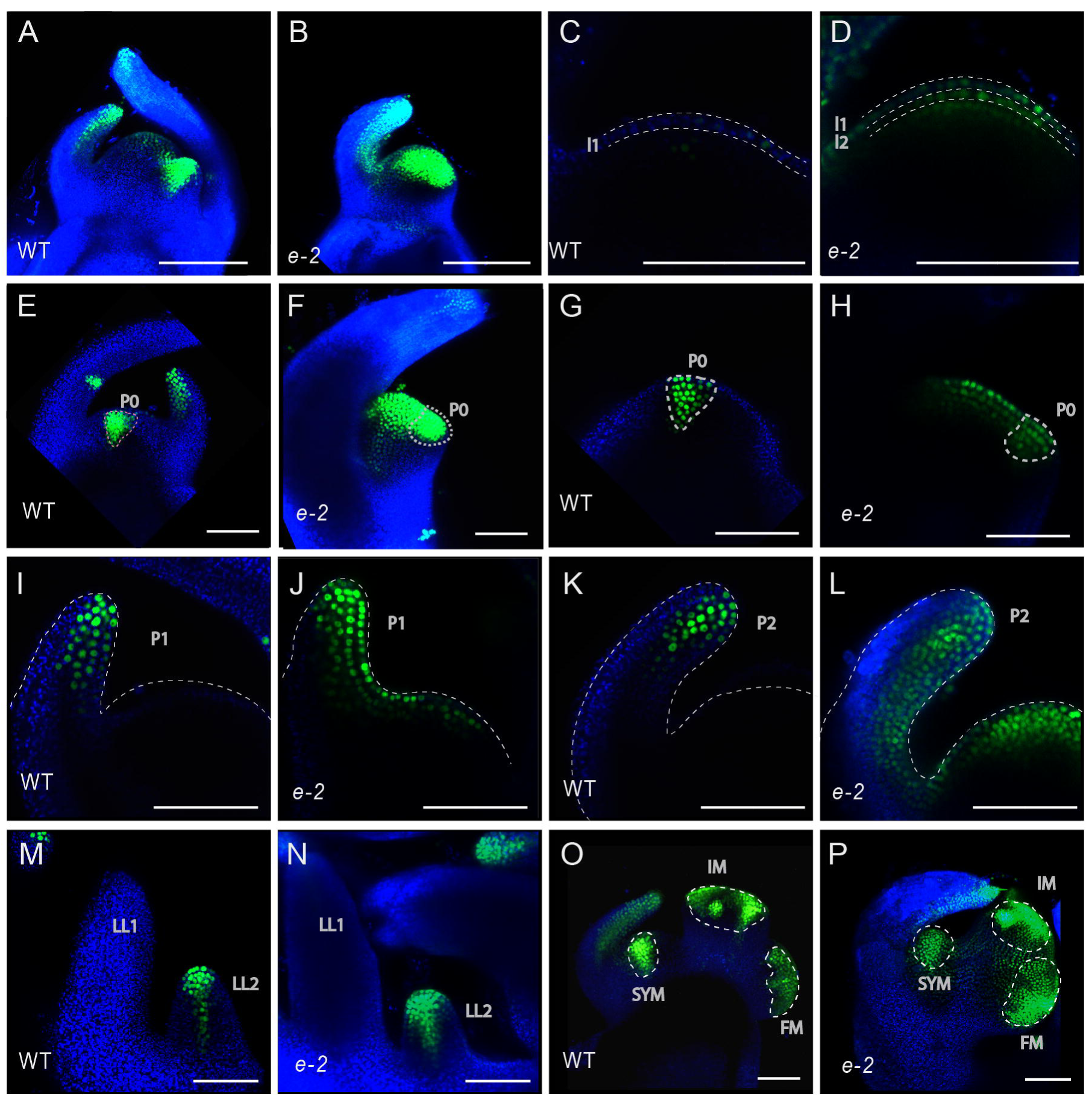
Auxin signaling in *e-2* mutant using DR5::Venus. Rendered z-stack of apical meristems (A) and (B) shows DR5::Venus (green) localization is expanded in (B) *e-2* compared to (A) wild type. Longitudinal section through the apical central zone shown in (A) and (B) revealing DR5::Venus expression in the first three layers of the apical meristem (L1, L2, and L3 layers) in (D) *e-2* and compared to (C) wild type, in which DR5::Venus expression is not visible. During leaf initiation (P0) in wild type (E) and (G) DR5::Venus expression is seen as a wedge shape pointing apically towards meristem center, while in *e-2* (F) and (H) DR5::Venus signal is more diffuse and does not penetrate as far inward. Longitudinal section through P1 (I) and (J) and P2 (K) and (L) show DR5::Venus signal is not separated from the DR5::Venus signal in the apical meristem of *e-2* (J) and (L) compared to wild type (I) and (K). During Lateral Leaflet (LL) initiation, DR5::Venus signal is wider in *e-2* (N) compared to wild type (M). (O) and (P) show DR5::Venus signal during sympodial inflorescence development. In wild type (O) DR5::Venus signal shows distinct separation between inflorescence meristem (IM), floral meristem (FM) and Sympodial Shoot Meristem (SYM). In *e-2* (P) the overall size is larger compared to wild type and DR5::Venus in continuous between all three (SYM, FM, IM) meristematic regions. Rendered images from z-stack (A), (B), (E), (F), and (M) to (P) while images from longitudinal sections are (C), (D), (G), (H), and (I) to (L). Scale bars = 100 μm.

In early developing leaf primordia *e-2* individuals show a similar pattern of DR5::Venus signal expansion that began in the incipient leaf, leaflet, and flower primordium. In wild type, DR5::Venus signal in P1-P3 developing organs and localizes throughout the center as the leaf develops, marking the site of the future midvein (Figure 6I and K). In *e-2*, DR5::Venus broadens into the adaxial side (Figure 6J) and internalized auxin transport through the center of the primordia does not narrow, creating a wider domain where the midvein will develop (Figure 6J and L). There is no separation of the DR5::Venus signal between early developing primordia and the apical meristem (Figure 6E and G); such a gap is normally seen in wild type (Figure 6J and L). Leaflets initiate similar to leaves, beginning as auxin maxima on the marginal blastozone of leaf primordia and continue with internal auxin transport through the center of the developing leaflet (Figure 6M) [6]. In *e-2*, there is more accumulation at the tip of the developing leaf primordia and the canalization of auxin through the center is widened (Figure 6N). During *e-2* inflorescence development the separation of auxin signaling between the IM and FM is not clearly delineated (Figure 6O). In addition, auxin maxima visualized by DR5::Venus as normally separate regions within the FM (Figure 6O) are often merged in *e-2* (Figure 6P). As interpreted by DR5::VENUS visualization, lack of *SlSoPIN1a* function in *e-2* causes a widening of the auxin flow pathway during leaf initiation, early leaf and sympodial meristem development, likely contributing to aberrant leaf morphology, vasculature, inflorescence, and flower defects found in *e-2* plants.

## Discussion

### e-2 is a *sopin1* mutant

In an attempt to identify *PIN1* and *SoPIN1* loss of function lines in *S. lycopersicum* we searched the TGRC mutant database for monogenic mutant lines previously mapped to the same chromosome of the three known *PIN1/SoPIN1* genes in *S. lycopersicum: SlSoPIN1a* (Solyc10g078370), *SlSoPIN1b* (Solyc10g080880), and *SlPIN1* (Solyc03g118740). We identified several lines that have abnormalities in leaf development and spatial patterning similar to other known *PIN1* loss of function mutants in *A. thaliana* and *Medicago truncatula* [26,45]. In the EMS induced mutant line *e-2 we* identified a C-T^490^ change, which causes a premature stop codon in the translated amino acid sequence of *SlSoPIN1a* (Figure 2A). C-T^490^ results in an inability to produce the SlSoPIN1a protein as evidenced by lack of SoPIN1a antibody signal in *e-2* apices (Figure 2E-I). Further evidence that C-T^490^ is responsible for the *e-2* phenotype comes from co-segregation and complementation analyses (Figure 2B and C). These studies verify that *e-2* is a loss of function mutant of *SlSoPIN1a*, a member of the *SoPIN1* clade [36-38] (Figure 1), sister to one of the most prolifically studied plant development genes, *PIN1*. We used detailed phenotypic characterization of the *e-2* mutant and visualization of the auxin inducible promoter DR5::Venus in this genetic background to develop an understanding of how *SlSoPIN1* functions in maintaining spatial patterning during early leaf and flower organogenesis in *S. lycopersicum*.

### *SlSoPIN1a* facilitates auxin localization in the L1 layer to establish phyllotaxy

Classically described by the Hofmeister rule, and based on the observation that leaves develop farthest from previous leaf initiation events, positioning of leaf primordia on the apical meristem was thought to be restricted by an inhibitory field around existing primordia [59]. It is now widely accepted that the interaction of *PIN1* and auxin is the leading mechanism underlying the Hofmeister spacing rule [4,26,27,56,60]. *PIN1* proteins direct auxin towards auxin maxima, draining auxin from surrounding cells and thus inhibiting the creation of new auxin maxima and new foci of leaf initiation nearby. Loss of *SlSoPIN1* function in *e-2/slsopin1* plants results in deviations from spiral phyllotactic patterning, including a consistent switch to distichous and decussate patterns (Figure 3A-E). Visualization of the auxin response reporter DR5::Venus reveals *e-2/slsopin1* apices have a dramatic expansion of the auxin response, including extension into L1, L2, and subepidermal layers throughout the entire dome of the apical meristem (Figure 3D-G). For establishing phyllotaxy, the importance of auxin transport in the L1 tissue layer has been proven extensively as 1. Models can accurately predict phyllotaxy while only incorporating information from the L1 layer [56,60]; 2. Laser ablation of the L1 layer results in auxin transport defects (Kuhlmeier); 3. *PIN1* expression in the L1 layer is sufficient to restore phyllotactic patterning in *atpin1* [61]. In addition knockout of *PIN1* in L1 and L2 abolishes lateral organ development, while knockout of *PIN1* in L1 only affects patterning [61]. Inferring auxin response as indicative of auxin localization, we conclude that *SoSlPIN1a* functions to regulate auxin distribution to the L1 layer, specifying convergences points that demarcate leaf initiation to establish phyllotactic patterning.

Convergence point formation also coincides with drainage of auxin from surrounding cells. The lack of *SoSlPIN1a* function in *e-2/soslpin1a* individuals likely causes build-up of auxin that results in drainage at multiple locations leading to aberrant broad internal auxin presence, explaining the subdermal DR5::Venus observed throughout *e-2/slsopin1a* apices (Figure 6D; Figure 7). The increase in auxin presence in *e-2/soslpin1a* meristems likely elicits auxin induced cell wall acidification, which activates expansin proteins [62] or pectin demethylesterification [63]. Meristem size may also contribute to changes in phyllotaxy observed in *e-2/soslpin1a* (Figure 3F-H) as suggested in previously characterized lines with phyllotactic patterning abnormalities [46-48]. The increase in meristem surface area could allow auxin maxima to form further away from previous initiation events, which is what we observe as divergence angles cluster around 180° (Figure 3F). Overall, the phyllotactic patterning observed in *e-2/soslpin1a* is likely the result of *SlSoPIN1a* mediated auxin transport being impaired in the L1 layer, limiting the ability to demarcate proper convergence points and the secondary result of changes in physical restraints caused from increase in meristem size.

**Figure 7.**
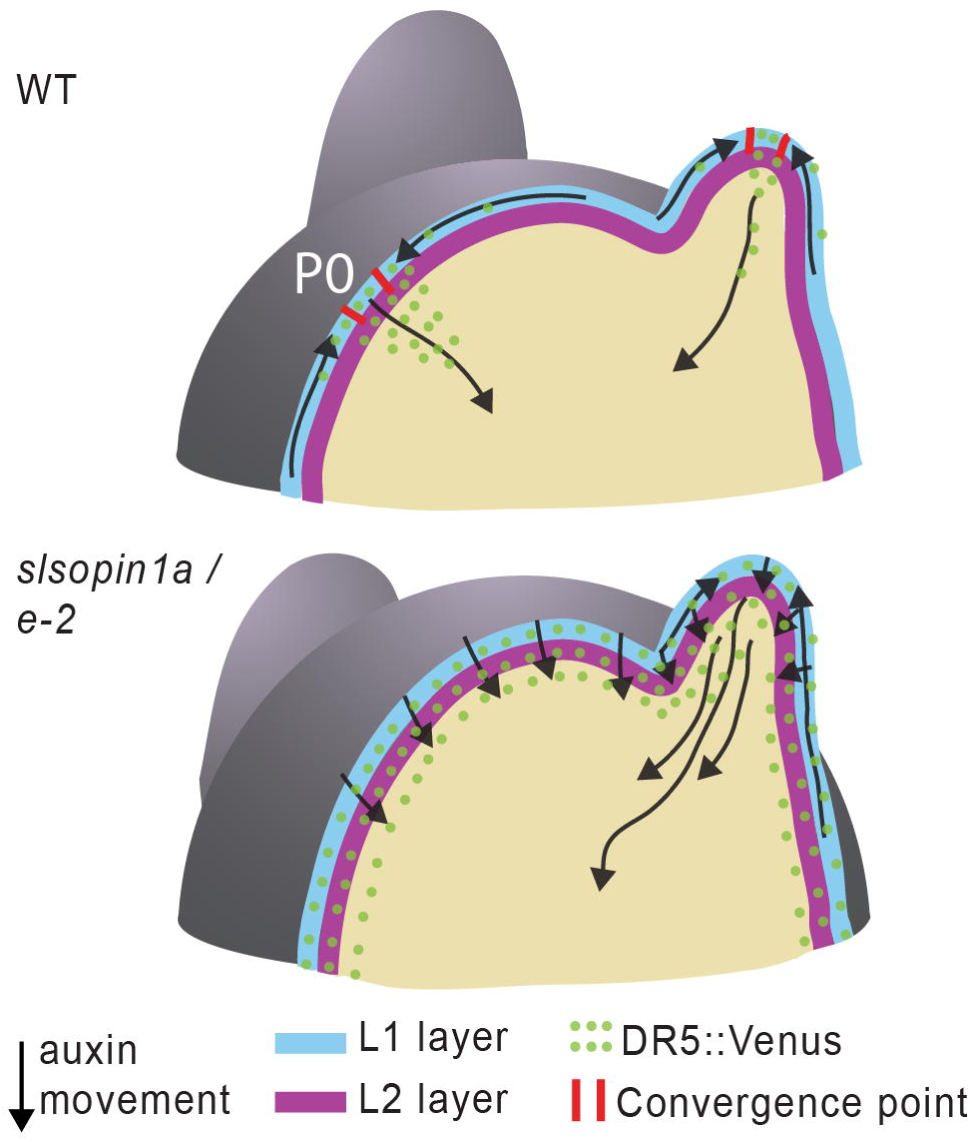
Schematic describing the role of auxin transport and localization in wild type and *e-2* apices. The top image represents a wild type meristem before sympodial growth. In wild type, auxin is transported along the L1 layer in both the meristem and newly initiating leaves. Auxin is transported to convergence points (red lines) which mark the site of leaf initiation (P0) on the meristem. At the convergence points there is the guidance of auxin basally guiding the development of veins. Emerging vein patterning occurs from the narrowing of canals of auxin transport. In *e-2*, auxin signaling (DR5::Venus) is found in the L1, L2, and subdermal layers of the meristem and newly developing leaf organs. Auxin transport is disturbed from lack of *SlSoPIN1a* function creating problems in establishment of convergence points, refinement and narrowing of vascular development, and leaf margin delimitation.

### *SlSoPIN1a* functions in spatial organization during leaf morphogenesis

Subsequent leaf morphogenetic processes also rely on the repeated pattern of *PIN1* convergence at the epidermal surface and internal auxin transport from convergence points, as seen during leaflet initiation and serration and lobe development [6,9,25]. In *e-2* individuals, leaflet initiation is compromised as complexity is overall decreased (Figure 4J) and leaflet spacing appears irregular compared to wild type (Figure 4I). DR5::Venus localization patterns during leaflet formation indicate a widening of auxin flux inwards as auxin canalization occurs (Figure 6I-N). This widening of auxin signaling and lack of refinement of internal auxin paths found in early developing *e-2/soslpin1a* leaf primordia likely contributes to the widened and fused vasculature observed in mature *e-2/soslpin1a* leaves (Figure 4C-H; Figure 7). Distortions in vascular development would also influence the spatial organization problems observed during later leaf development in *e-2/soslpin1a* mutants. Leaf morphogenesis appears to worsen through time as exemplified by laminal bulging seen in *e-2/slpin1a* leaves, which become increasingly rugose with developmental age (Figure 4A and B; S3 Figure A-E). The cumulative result of the many early problems in auxin directed leaf development in *e-2/soslpin1* plants can be seen in the final leaflet shape which is extremely varied (Figure 4K). In addition, placement of serrations and lobing in *e-2* is more random and asymmetric, resulting in an average outline shape lacking lobes all together (Figure K and M). The importance of *SoSlPIN1* in both leaflet formation and leaf initiation is clear and the gene appears to function in directing auxin to specify proper placement of leaf developmental features. Since most leaf developmental processes are reiterative and self directing, the aggregate effect of misplaced auxin at leaf initiation shows cumulative results in the extremely aberrant final leaf phenotype seen in *entire-2/slsopin1a* plants.

### *SlPIN1a* functions in meristem maturation and organ specification

Of all the *PINs, PIN1* appears to be the most important for reproductive development [3,13,26,64]. In *S. lycopersicum, SoSlPIN1a* functions in both the positioning and specification of lateral organs during sympodial shoot meristem and inflorescence development. The sympodial unit on the main axis in wild type plants consists of one complex leaf and an inflorescence, while *e-2/soslpin1a* possesses dramatic variation in number of inflorescences and leaf number per sympodial unit (Figure 5A). The inflorescence unit of *e-2/soslpin1a* bears both flowers and leaf organs (Figure 5C-J), which is unusual, as all varieties of *S. lycopersicum* bear inflorescences with only flower units (Figure 5B and J) [53]. Meristems are thought to establish reproductive identity through a defined maturation process, in which the likelihood to produce leaf organs, often defined as “vegetativeness”, decreases over time and ends with the determinate floral meristem identity [65]. The inflorescence in *e-2/soslpin1a* may mature slower than wild type and possess a higher degree of vegetativeness. This slowing of maturation could result in a maturation state similar to sympodial growth on the main axis (Figure 5A), which would explain the presence of leaf-like organs in *e-2/slsopin 1a* (Figure 5B-L). This hypothesis would explain the occasional presence of complex leaves, which closely resemble main axis leaves on the inflorescence axis (S4 Figure C).

An alternative interpretation of the *e-2/soslpin1* inflorescence phenotype is that the leaf organs found on the inflorescence are bracts - leaf like organs that subtend flowers. Wild type *S. lycopersicum* plants do not contain visible bracts, but in the Solanaceae reproductive branching systems manifest in a variety of ways [53]. In many green-fruited wild relatives of *S. lycopersicum*, the first determinate organ in a floral branching system is a leaf-like bract [54,66]. It has even been suggested that all Solanaceae inflorescences have bracts, but their development may have been suppressed early [67], as seen in *A. thaliana [68-71]* and potentially in *S. lycopersicum [72]*. Modulation of auxin localization may be an evolutionary strategy for transformation of inflorescence types. The phenotype of leaf-like organs found on *e-2/slsopin1a* inflorescences could be a modulation in inflorescence branching identity, allowing the development of visible bract units as seen in closely related wild species. The most recent *SoPIN1* duplication event in *S. lycopersicum* occurred sometime before Solanaceae radiation (Figure 1B). It would be interesting to test if modulation of *PIN1* and *SoPIN1* gene expression regulates inflorescence branching identity in diverse range of species in Solanaceae. Recent gene duplication events in the *PIN1* clade are common across angiosperms, as revealed through phylogenetic analyses clades (Figure 1A and B) [36,37]. These recently evolved paralogs may represent an evolutionary mechanism to finely tune auxin directed development. Only more functional work in *PIN1* and *SoPIN1* within closely related species can reveal evolutionary functional divergence in the larger *PIN1* clade.

### Subfunctionalization and functional redundancy of *PIN1* and *SoPIN1* genes across species

Our current understanding of *PIN1* directed auxin transport has largely ignored the contribution of *SoPIN1* genes because the most intensely studied species, *A. thaliana*, has lost the *SoPIN1* clade (Figure 1B) [36,37]. Most angiosperm species have at least one representative in each of the *SoPIN1* and *PIN1* clades (Figure 1B) [36,37]. This level of conservation throughout angiosperm species suggests both *SoPIN1* and *PIN1* genes have conserved function in plant development. Evidence that *AtPIN1* and *SoSlPIN1* share function is that both AtPIN1 and SoSlPIN1 proteins show very similar expression patterns, localizing predominantly in the L1 layer and subepidermally directing auxin during leaf initiation events (Figure 2E, Figure S3 N-U) [39]. Further support of shared function is results from our complementation experiment in which *AtPIN1* is capable of rescue of the *e-2* spiral phyllotaxy (Figure 2C and E). Even though *AtPIN1* and *SlSoPIN1a* likely share similar roles, the loss of function phenotype in both species is very different, implying functional divergence. Loss of *AtPIN1* function in *A. thaliana* results in loss of lateral organ initiation after flowering, while in *e-2 /slsopin1a*, subdermal auxin flux is delayed and *e-2/slsopin1a* initiates leaves at a slower rate than wild type (S3 Figure M) and proceeds with subsequent lateral organ initiation events throughout the lifespan of *e-2/slsopin1a* plants. A likely reason for the extreme phenotype of the *atpin1* mutant is that *AtPIN1* holds the function of both the *SoPIN1* and *PIN1* clades resulting from the recent loss from any *SoPIN1* representative in *Brassicacea* (Figure 1) [36,37].

With recent research in organisms outside *A. thaliana* we have begun to uncover the possible evolutionary consequences of subfunctionalization of *PIN1* and *SoPIN1* genes. The *SoPIN1* and *PIN1* gene have distinct functions at the tissue layer level, as seen in the separation of *PIN* function during root development [73]. Duplication and subfunctionalization of the ancestral *PIN1* gene led to an uncoupling of auxin transport, resulting in distinct functions assigned to *SoPIN1a* and *PIN1* genes in the role of L1 auxin transport to create convergence points, subepidermal auxin flow, and the narrowing of auxin flux channels to refine emerging veins. Recent work in *Brachypodium distachyon* (Brachypodium) found that *Brachypodium SoPIN1 (BdSoPIN1)* gene expression localizes predominantly in the L1 layer while the two *Brachypodium PIN1* genes *(BdPIN1a* and *BdPIN1b)* are expressed subepidermally. Through this work the authors conclude that in *Brachpodium SoPIN1* functions in marking the sites of organ formation, while the *BdPIN1a* and *BdPIN1b* function in internalization of auxin flow to direct vascular development [37]. Our work does not support such a distinct separation, but DR5::Venus localization differences found in *e-2* (Figure 6) suggest *SoPIN1* functions predominantly in L1 auxin transport, it also contributes to canalization processes as evidenced by the vascular phenotypes found in *e-2* leaves (Figure 4C-H).

While it appears that subfunctionalization of *PIN1* and *SoPIN1* genes play a role in auxin distribution at the tissue level [37], another important level for morphogenetic outcomes is that of developmental timing. As discerned from studies on the *slm-1 (PIN 10/ Medtr7g*1*06430)* mutant, *Medicago truncatula SoPIN1* function is vital during juvenile leaf initiation; the juvenile leaf is completely abolished in *slm-1* and later leaf development has problems similar to those observed in *e-2/slsospin1a* [45]. In *A. thaliana*, leaf development can be split into three developmental stages which are morphologically distinct [27]. In *atpin1*, the leaf phenotype clearly varies in the three leaf stages[70], as leaf vasculature and shape gets progressively impaired with age, until there is a complete loss of lateral organ development after bolting [27]. There are two developmental stages in *S. lycopersicum* leaf formation, juvenile leaves, and adult leaves before and after sympodial growth. In *e-2/soslpin 1a* phyllotaxy abnormalities occur during juvenile development (Figure 3A-E), and developmental problems prior to sympodial growth are observed, but the specification of organ type (flower vs leaf) remains identical to wild type. After *e-2/soslpin1a* begin sympodial growth, specification of organ identity becomes extremely aberrant (Figure 5A-L). Thus, auxin flux as regulated by *SoSlPIN1a* is involved in organ identity throughout developmental age. Further, while *SoPIN1* gene expression remains constant during the early stages of monopodial growth, expression levels change during sympodial meristem development both in time and at the tissue level to specify inflorescence identity (S7 Figure). Therefore, a uniting feature of all *PIN1* and *SoPIN1* loss of function mutants is the insight they provide into the role of auxin through developmental time.

## Conclusion

*PIN1* directed auxin transport during early shoot organogenesis regulates the reiterative process of shoot development. The repeated developmental module of epidermal auxin transport and internal auxin transport is recycled during margin growth, leaf and leaflet formation, and positioning of vasculature. The importance of PIN1 in directing these processes is undisputed, yet it is still unknown how this mechanism has evolved through time. The extent of partitioning of auxin transport functions within *PIN1* and *SoPIN1* clades, and the functional consequences of this partitioning may provide explanations for the immense diversity in leaf form and inflorescence architecture seen in nature. The advent of available sequencing technologies for genome-scale gene identification and ability to undertake functional studies in a multitude of species has set the stage for comparative studies of developmental processes across Angiosperm clades. These studies should help uncover how the *PIN1* and *SoPIN1* clades contributes to the astounding morphological diversity found in the plant kingdom.

## Materials and Methods

### Phylogenetic analysis

We used only sequences from species from fully sequenced genomes to ensure proper representation of the PIN gene family. All cDNA sequences were retrieved from either Pytozome [74], or through BLAST searches using genome databases available on the Sol Genomics Network [75]. Gene orthology was further confirmed through comparisons with previous phylogenetic work [36,37]. *AtPIN3* (AT1g70940) was used as the outgroup. All sequences were aligned using MUltiple Sequence Comparison by Log-Expectation (MUSCLE) alignment [76] on the EMBL-EBI bioinformatics web server [77]. Sequences were further trimmed using TrimAL (version 1.2rev59) with gap threshold set to 90% and a specified conservation minimum of 60% positions from the original alignment. Using the servers from The CIPRES Science Gateway (version 3.3; [78]. Maximum likelihood analysis was performed using RAxML-HPC2 (version 8.1.24; [79]. Visualization and editing of trees was accomplished using FigTree (version 1.4.2; [80]. Maximum Liklihood tree and alignment can be found at Treebase: http://purl.org/phylo/treebase/phylows/study/TB2:S18794?x-access-code=e8166b52ee913f392b9454be09ef962d&format=html.

### Plant material and growth conditions

Seeds of *e-2* (accession 3-705) and control (LA3130) were obtained from the Tomato Genetics Resource Center (TGRC). The transgenic DR5::Venus *(cv M82)* lines were previously described by and AtpPIN1::PIN1::GFP *(cv Moneymaker)* [39]. To ensure developmental synchronization, all seed lines were first sterilized with 50% bleach for 2 minutes, rinsed 10x with distilled water, and then placed on a moist paper towel in Phytotrays (Sigma-Aldrich) under dark conditions for two days before being placed in a growth chamber (temperature 22 **°**C, 16:8 light-dark cycle) for three days before transplanting to soil.

### Microscopy

Microscopy of apices was performed using a Zeiss Discovery V12 stereomicroscope and photographed using an AxioCam MRc digital camera (Carl Zeiss MicroImaging, Thornwood, NY, USA). For fluorescent imaging, the microscope was equipped with an X-Cite 120 light source, a pentafluor GFP wideband cube (Zeiss KSC 295-831D, excitation HQ 470/440 nm and dichroic mirror 495LP) and a long-pass emission filter (KS295-831WD, 500 nm). Images for immunolocalization and histology images were captured using a Nikon Eclipse E600 compound microscope and a Nikon digital camera (Nikon, Melville, New York, USA). Some photographs were adjusted for brightness and contrast and assembled into figures using Adobe Photoshop CS6 and Adobe Illustrator CS6 (Adobe Systems, San Jose, CA, USA).

### Measurement of angle divergence and meristem size

Measurement of angles were made on plant apices of both wild type (LA3130) and *e-2* were harvested 20 days after germination. All measurements were made on leaves before sympodial growth. The apices were then fixed in 3:1 acetic acid:ethanol and embedded in 100% paraffin. The apices were sectioned onto slides and stained with toluidine blue. Sections were visualized and photographed using microscopy methods as described above. In addition, older plants (50 days after germination) were also measured. In the older plant population, the leaflets were removed, leaving only the petiole, these were labeled and aerial photographed above the plant were taken using Olympus SP-500 UZ camera. ImageJ (version 1.46R) was then used to calculate angle divergence of the first five to eight leaves. For quantifying meristem size, 12 day old plants were dissected to when meristem was visible and then photographed under a dissecting microscope at 100x magnification. The meristems were measured in ImageJ (version 1.46R) using the straight line tool. Analysis and visualization was performed in using R [81] using R package ggplot2 [82]. Analysis scripts are available at https://github.com/iamciera/sister-of-pin1-material.

### Leaf analysis: shape, complexity, and clearing

Plants were harvested 51 and 52 days after sowing, having six fully expanded leaves and were grown in a walk-in chamber (Conviron), temperature 22 C, 16:8 light-dark cycle. Leaflets were removed from the petiole of L1-L7 and placed under non-reflective glass. Cameras (Olympus SP-500 UZ) were mounted using a Adorma; 36’ Deluxe Copy Stand, and remotely controlled with Cam2Com software (Sabsik). Normalization was made from measurements from rulers present in each photograph. Shape analysis was accomplished using ImageJ, by converting the photographs to binary images, with subsequent measurements of area, perimeter, circularity, aspect ratio, roundness, and solidity. Complexity was measured by counting all leaflets present on each leaf. The binary images were then processed using the Momocs package in R [51] to determine the elliptical fourier descriptors, analysis script is available at https://github.com/iamciera/sister-of-pin1-material.

### Immunolocalization

Apices of 14 day old plants were fixed and vacuum infiltrated in 3:1 Methonal: acetic acid. Tissue then went through an ethanol series (10, 30, 50, 70, 85 2x100%) for thirty minutes each step mixed with PBS. Tissue was then incubated ethonal:PEG1500 (Sigma-Aldrich) series (3:1, 1:1, 1:3, 2x 1:1) at 45 **°**C. Tissue was embedded and mounted on Saline Prep slides (Sigma-Aldrich) following tissue transfer methods previously described in Gao and Godkin, 1991[83]. Blocking, secondary, and primary anti-body incubation was performed as previously described in [84]. Primary SlSoPIN1a antibodies were generated as described in [39]. SlSoPIN1a antibodies were diluted 1:200 and secondary antibody (Alexa Fluor 488-conjugated goat anti-rabbit) was diluted 1:300. Control slides, which were incubated with only secondary antibodies, were included in all experiments.

## Supplemental Information

**S1 Figure - Phenotypic characterization of *S. lycopersicum* lines from the TGRC mutant database**. (A) to (G) Images of *S. lycopersicum* lines that have been roughly mapped close to known *SlPIN1, SlSoPIN1a*, and *SlSoPIN1b* genes and the wild type backgrounds. Alisa Craig (AC) (A) is the wild type background for *solanifolia (sf)* (B), *restricta (res)(C), divaricata (div)* (B), and *oivacea (oli)* (E), while 3130 (F) is the wild type background for *entire-2 (e-2)* (*G*). (H) and (I) are images of Leaf 4 from four week old plants. Scale bars = 20 mm.

**S2 Figure - Confirmation analysis to verify *entire-2* is a *SlSoPIN1a* loss of function mutant**. (A) wild type and (B) *e-2* plants at 40 days old (C) RT-PCR showing no difference in transcript between wild type and *e-2* in *SlPIN1* (blue), *SoSlPIN1a* (green), and *SoSlPIN1b* (pink) from four biological replicates. (D) Bar graph illustrating that plants that have been complemented with PIN1∷GFP have higher leaf complexity as measured by counting the total number of leaflet number on Leaf 1-4 of four week old plants. (E) Difference in divergence angle seen across the first seven leaves in wild type, *e-2*, and AtPIN1∷PIN1∷PIN1 complemented individuals, illustrating that in wild type and complemented plant divergence angles cluster around 137.5**°**, while *e-2* plants tend to alternate from above and below 137.5**°** (as seen in distichous and decussate branching patterns).

**S3 Figure - pPIN1∷PIN1∷GFP and SlSoPIN1a protein localization in *S. lycopersicum apices***. (A) to (H) pPIN1∷PIN1∷GFP localization in wild type (A) to (D) and *e-2* (E) to (H) apices. pPIN1∷PIN1∷GFP localization pattern does not differ between the two lines during meristem (B) and (F), P1 (C) and (G), and P3 (D) and (H) development, suggesting pPIN1∷PIN1∷GFP complements by establishing proper functional PIN1 localization in *e-2*. (I) to (P) Immunolocalization using an antibody raised against SlSoPIN1a protein sequence in wild type apices. Protein localization was found during leaf initiation on the meristem (I) and (K), vasculature initiation (J), (L), and (N), and axillary meristem establishment (O) and (P) as previously reported [39].

**S4 Figure - *SlSoPIN1a* loss of function leads to developmental defects during *S. lycopersicum* leaf morphogenesis**. (A) Images of terminal leaflets of Leaf 4 from wild type and *e-2*, showing multiple midveins in the *e-2*. (B) to (E) Laminal (blade) tissue in wild type and and *e-2* terminal leaflets. Images of laminal bulging present in *e-2* leaves (C) and (E), which becomes worse as leaf development proceeds (compare *e-2* Leaf 4 (C) with leaf 5 (E). (F) to (K) Transverse sections stained with toludine blue of petiole (F) to (I) and leaflet midvein (J) to (K) showing vasculature fusing in *e-2* (G), (I), and (K) compared to wild type (F), (H), and (J). (L) to (O) Images of cotyledon phenotypes found in wild type (L) and *e-2* (M) to (O). Ratio represents frequency of phenotype found in wild type (n = 41) and *e-2* (n = 43) populations. (P) Box plot illustrating the number of leaves greater than 5 mm at 30 days after germination in wild type (grey) and *e-2* (blue) showing *e-2* has less leaves suggesting *e-2* develops at a slower rate than wild type. (Q) and (R) Scatter plots of Principal Components (PC) obtained from elliptical fourier analysis performed on leaf outlines from wild type (black) and *e-2* (blue). (Q) PC1 vs PC2 (R) PC3 vs PC4. Scale bars = 1mm in (B) to (K), 50mm in (D) and (E), and 10mm in (L) to (O).

**S5 Figure - The *entire-2* mutant of *S. lycopersicum* shows developmental defects during flower development** (A) to (C) Images of sepal from wild type and *e-2*. Sepals are larger in plants heterozygous for the C-T^490^ (+/-) (B) and to a greater extent homozygous (*e-2*) (C) plants compared to wild type (A). (D) and (E) inflorescences of *e-2* have presence of fully expanded simple (D) and complex leaves (E). (F) and (G) Images of young inflorescences of wild type and *e-2*. Leaf-like organs are present early in *e-2* inflorescence development (G). (H) and (I) Close up image of a leaf-like organ from an *e-2* inflorescence. Scale bars = 10mm in (A), (B), and (C), 4mm in (F) and (G), and 1mm in (H) and (I).

**S6 Figure - Characterization of fruit development in *entire-2***. Ovaries prior to fertilization from wild type (A) and (B), heterozygous (+/-) (E) and (F), and *e-2* (I) and (J). Developing fruit three days after fertilization in wild type (C) and (D), +/- (G) and (H), and *e-2* (K) and (L). Fruit prior to ripening (M) and (N) and after ripening (O) and (P). Scale bars = 1mm in (A) to (L), 50mm in (M) to (P).

**S1 Table - Phenotypic characterization of wild type and *entire-2* terminal leaflets**. Measurements were made on binary images made from terminal leaflets of leaves 1 - 4 of six week old plants. Numbers presented as mean ± standard deviation. * indicates the differences between wild type and *e-2* are significant at P < 0.05. Circularity score of 1 signifies a perfect circle, while the closer to zero, the leaves are progressively more lobed. All measurements were made in ImageJ.

**S2 Table - Measurements of meristem size in wild type and *entire-2***. Meristem size of 12 day old plants. Numbers presented as mean ± standard deviation. * indicates the differences between wild type and *e-2* are significant at P < 0.05.

## Author Contributions

Conceived and designed the experiments: CM, NS, DK, DC. Performed the experiments: CM, DK, DC. Analyzed the data: CM, NS, DK, DC. Wrote the paper: CM, NS. Edited the paper: CM, NS, DK, DC.

## Acknowledgements

The UC Davis Tomato Genetics Resource Center provided tomato germplasm, AtpPIN1∷PIN1∷GFP and the DR5:VENUSx6 lines are gifts from Cris Kuhlemeier (University of Bern) and Naomi Ori (Hebrew University, Israel), respectively. We would like to thank Paradee Thammapichai, Jeanice Jones, Sharon Zimmerman, Gillie Ish-Shalom, and Surbhi Chophla for technical assistance. Siobhan Braybrook generously provided the immunolocalization protocol. Thanks also to Siobhan Brady and Andrew Groover for their helpful comments. C.C.M was supported by a National Science Foundation Graduate Research Fellowship (DGE-1148897), Katherine Esau Summer Fellowship, Walter R. and Roselinde H. Russell Fellowship, and Elsie Taylor Stocking Fellowship. Part of the work was supported by NSF PGRP grant IOS-0820854 (to N.R.S., Julin Maloof and Jie Peng). This work used computational resources or cyber-infrastructure provided by the iPlant Collaborative (www.iplantcollaborative.org). The iPlant Collaborative is funded by National Science Foundation Grant DBI-0735191.

